# Decomposing cell identity for transfer learning across cellular measurements, platforms, tissues, and species

**DOI:** 10.1101/395004

**Authors:** Genevieve L. Stein-O’Brien, Brian S. Clark, Thomas Sherman, Cristina Zibetti, Qiwen Hu, Rachel Sealfon, Sheng Liu, Jiang Qian, Carlo Colantuoni, Seth Blackshaw, Loyal A. Goff, Elana J. Fertig

## Abstract

New approaches are urgently needed to glean biological insights from the vast amounts of single cell RNA sequencing (scRNA-Seq) data now being generated. To this end, we propose that cell identity should map to a reduced set of factors which will describe both exclusive and shared biology of individual cells, and that the dimensions which contain these factors reflect biologically meaningful relationships across different platforms, tissues and species. To find a robust set of dependent factors in large-scale scRNA- Seq data, we developed a Bayesian non-negative matrix factorization (NMF) algorithm, scCoGAPS. Application of scCoGAPS to scRNA-Seq data obtained over the course of mouse retinal development identified gene expression signatures for factors associated with specific cell types and continuous biological processes. To test whether these signatures are shared across diverse cellular contexts, we developed projectR to map biologically disparate datasets into the factors learned by scCoGAPS. Because projecting these dimensions preserve relative distances between samples, biologically meaningful relationships/factors will stratify new data consistent with their underlying processes, allowing labels or information from one dataset to be used for annotation of the other—a machine learning concept called transfer learning. Using projectR, data from multiple datasets was used to annotate latent spaces and reveal novel parallels between developmental programs in other tissues, species and cellular assays. Using this approach we are able to transfer cell type and state designations across datasets to rapidly annotate cellular features in a new dataset without a priori knowledge of their type, identify a species-specific signature of microglial cells, and identify a previously undescribed subpopulation of neurosecretory cells within the lung. Together, these algorithms define biologically meaningful dimensions of cellular identity, state, and trajectories that persist across technologies, molecular features, and species.

**GRAPHICAL ABSTRACT:** 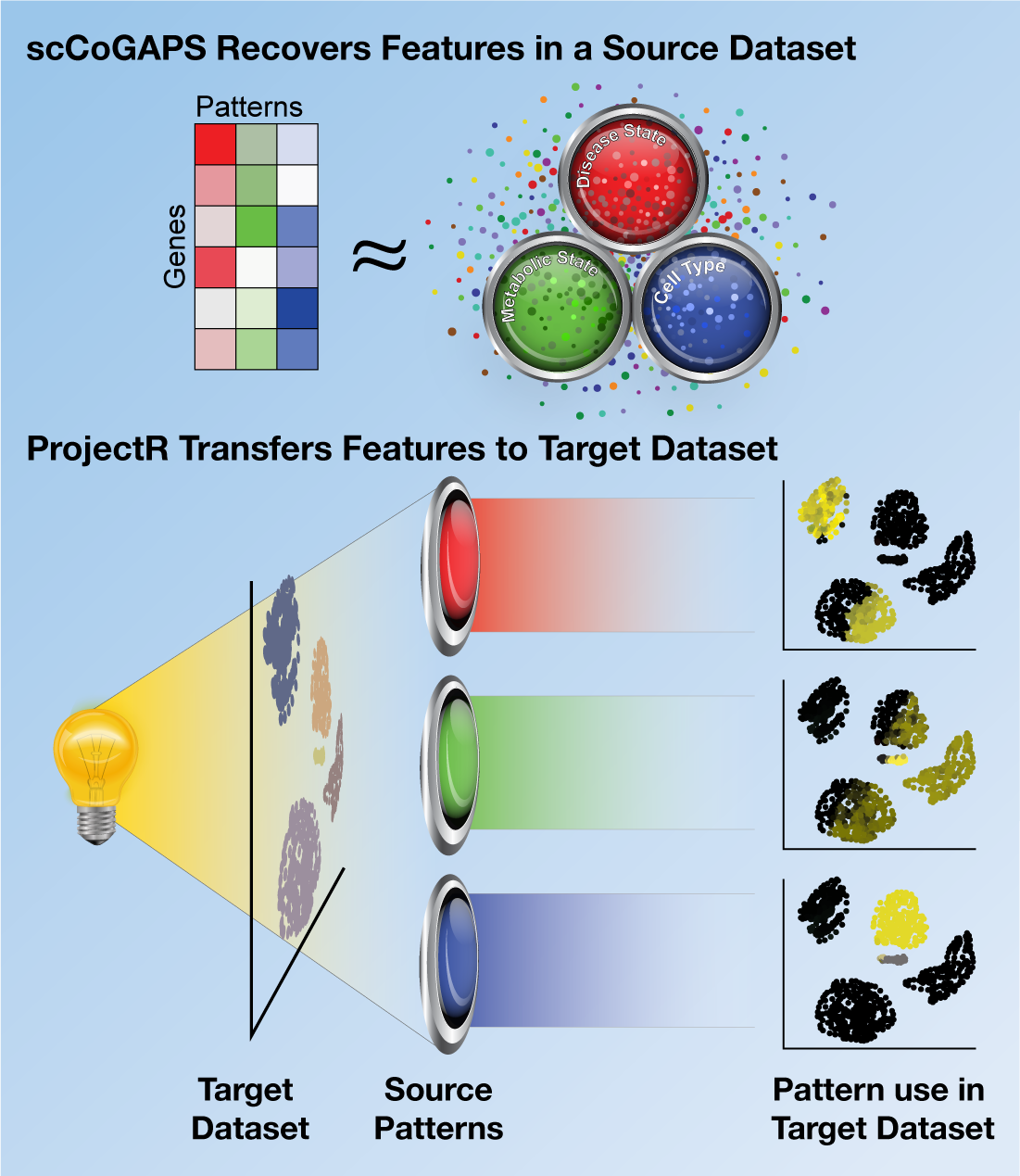

## INTRODUCTION

The transcriptional identity of an individual cell is determined by the combinatorial effects of diverse biological processes. The additive effects of these processes on transcriptional regulation, mRNA processing, and mRNA degradation are encoded in a cell’s gene expression profile. Dimension reduction techniques deconvolve gene expression data into discrete latent spaces that comprise signatures of these biological processes (Brunet et al., 2004; Cleary et al., 2017a; Kossenkov et al., 2007; Stein-O’Brien et al., 2017a; Wagner et al., 2016; Zhu et al., 2017). This decomposition of the transcriptional profile into signatures of individual processes enables a biologically meaningful interpretation of cellular activity arising as the additive contribution of multiple concurrent systems within the cell, each with their own effects on the transcriptome. In this manner, individual patterns, corresponding to both biological and technical influences on the transcriptome, can be construed as constitutive components of the cellular steady state, and independently examined or compared across conditions (Wagner et al., 2016).

Single cell analysis of gene expression provides an ideal approach to learn these signatures, since independent measures correspond to the ensemble of processes in an individual cell, and are not confounded by the effects of aggregating functionally distinct cell types and states. While scRNA-Seq is poised to revolutionize our understanding of cell type classification, it is critical to first develop an appropriate representation of the patterns of gene expression that generate the unique transcriptional signatures of an individual cell. This representation is especially critical in the mammalian central nervous system (CNS), which consists of thousands of related but functionally distinct cell types. The hierarchical nature of CNS development further obscures clear distinctions between bona fide cell types, cell subtypes, and physiological or random variation in gene expression within individual subtypes (Tasic et al., 2016).

Latent space techniques perform dimension reduction to identify pertinent biological processes in scRNA-Seq data. These techniques identify a subset of all measured genes that are expressed to different degrees depending both on the identity of, and active biological processes in each cell. In this approach, each learned space is called a factor or pattern with a corresponding amplitude or weight for each feature (gene). A single gene may be expressed in multiple cell types and states, but the patterns of expression within the set of genes are signatures of cellular identity and state (Cleary et al., 2017b). Numerous latent space techniques are emerging to learn these low dimensional gene expression patterns from scRNA-Seq data (Stein-O’Brien et al., 2017a; Wagner et al., 2016; Wu et al., 2017). These dimension reduction techniques have had similar success in deconvolving biological processes arising between mixtures of cell types in bulk data (Stein-O’Brien et al., 2017a). In both cases, complementary, biologically-driven assessment tools are critical to determine whether the feature representations in the low-dimensional latent space reveal gene weights that reflect the basal processes of cells.

One method to determine the biological relevance of low dimensional representations of cellular processes is to determine their robustness in data from multiple sources and contexts. For example, factors that are associated with related biological processes in multiple datasets can be compared (DeTomaso and Yosef, 2016; Kiselev et al., 2018). However, differences in information content across platforms and technical artifacts including batch effects (Hicks et al., 2017; Leek et al., 2012), library preparation biases (Li and Ngom, 2013), and antibody quality, can dominate signals in multi-platform studies, challenging such direct comparisons. These differences are all the more pronounced in datasets from different biological conditions. Instead, classifiers can be designed to relate predicted cellular identity or biological processes in new target datasets based upon the latent spaces learned in a source dataset. Specifically, biologically meaningful relationships/features will stratify new data consistent with their underlying biological processes, while dataset-specific factors such as those associated with batch effects, will result in little to no information content in the projection as they are by definition, unique signatures of a dataset.

Transfer learning (TL) techniques use latent space factors learned from one or more sources to improve learning of a new target dataset. TL does not require that source and target data have the same distribution, domain, or feature space (Pan et al., 2008; Torrey and Shavlik, 2009). Instead, TL exploits the fact that if two datasets share common biological processes, a feature mapping between the two datasets can be used to identify and characterize relationships between the data defined by individual latent spaces. (Pan et al., 2008). In the genomics space, a natural feature mapping exists as comparable gene features between source and target datasets, including orthologous mapping of genes across species. Furthermore, TL is able to use component measures learned via latent space techniques to bridge high-dimensional datasets. As a result, TL methods are particularly well suited for integrating multi-omic analyses across data platforms, modalities, and studies.

To adapt TL approaches for use in high-throughput genomic data analysis, we propose that low dimensional representations from latent space analyses of transcriptional data provides a natural, continuous representation of biological or technical processes encoded in the gene expression profile of a cell. We further hypothesize that these low dimensional representations can be used as a lens through which to view the usage of a specific process or feature across multiple conditions. To test this conjecture, we have developed three theoretical advances: (1) a non-negative matrix factorization (NMF) technique, scCoGAPS, that embeds the technical and biological structure of scRNA-Seq data in latent space discovery, (2) a scalable computing framework for latent space discovery and cross-validation of factors, and (3) a transfer learning technique, projectR, to query the relationship of latent spaces learned in one biological context against the biological processes that occur in another.

Knowledge transfer via projectR was designed with speed and scalability in mind. ProjectR’s ability to rapidly transfer annotation, classify cells, and identify the use of biological processes greatly accelerates the rate of biological discovery in high dimensional data when used in conjunction with manual annotation and expert curation. In line with the fundamental tenet of TL, projectR is agnostic to a priori knowledge or annotation within the target data. The ability to rapidly test relationships and transfer knowledge via projectR demonstrate the potential of TL as a powerful tool for *in silico* experimentation and hypothesis generation. While we focus this application of projectR to low dimensional factors learned with scCoGAPS, the algorithm generalizes as an exploratory analysis and biological interpretation method for other dimension reduction, or latent space discovery techniques as well.

We complement the methods developed in this manuscript with application to a time course scRNA-seq dataset from murine retina development. Application of scCoGAPS to these data identified gene expression signatures of discrete cell types, as well as cellular processes controlling neurogenesis and cell fate specification. We then applied projectR to transfer knowledge described by these dimensions into multiple target datasets including bulk RNA-Seq, ATAC-Seq and scRNA-Seq. This TL method is able to annotate latent spaces and reveal novel parallels between the scCoGAPS retinal development patterns and gene expression patterns observed in different tissues, molecular features, and species. Together, these algorithms define biologically meaningful dimensions of cells identity, state, and trajectories that persist across platforms and species.

## RESULTS

### Adaptive sparsity for learning factors from scRNA-Seq (scCoGAPS): Theory

ScCoGAPS is an NMF algorithm. NMF algorithms factor a data matrix into two related matrices containing gene weights, the Amplitude (**A**) matrix, and sample weights, the Pattern (**P**) Matrix (Fig 1A). In this case, we assume that the data matrix is log-transformed fragments per kilobase of RNA per million mapped reads (FPKM) expression estimates in the case of bulk RNA-seq data and log-transformed unique molecular identifier (UMI) in the case of 10x Genomics 3’-end tagging scRNA-Seq data. Each column of **A** or row of **P** defines a factor, and together these sets of factors define the latent spaces among genes and samples, respectively. Each sample-level relationship in a row of the pattern matrix is also referred to as a pattern and the corresponding gene weights the amplitudes. In NMF, the values of the elements in the **A** and **P** matrices are constrained to be greater than or equal to zero. This constraint simultaneously reflects the non-negative nature of gene expression data and enforces the additive nature of the resulting factors, generating solutions that are biologically intuitive (Lee and Seung, 1999).

**Figure 1.**
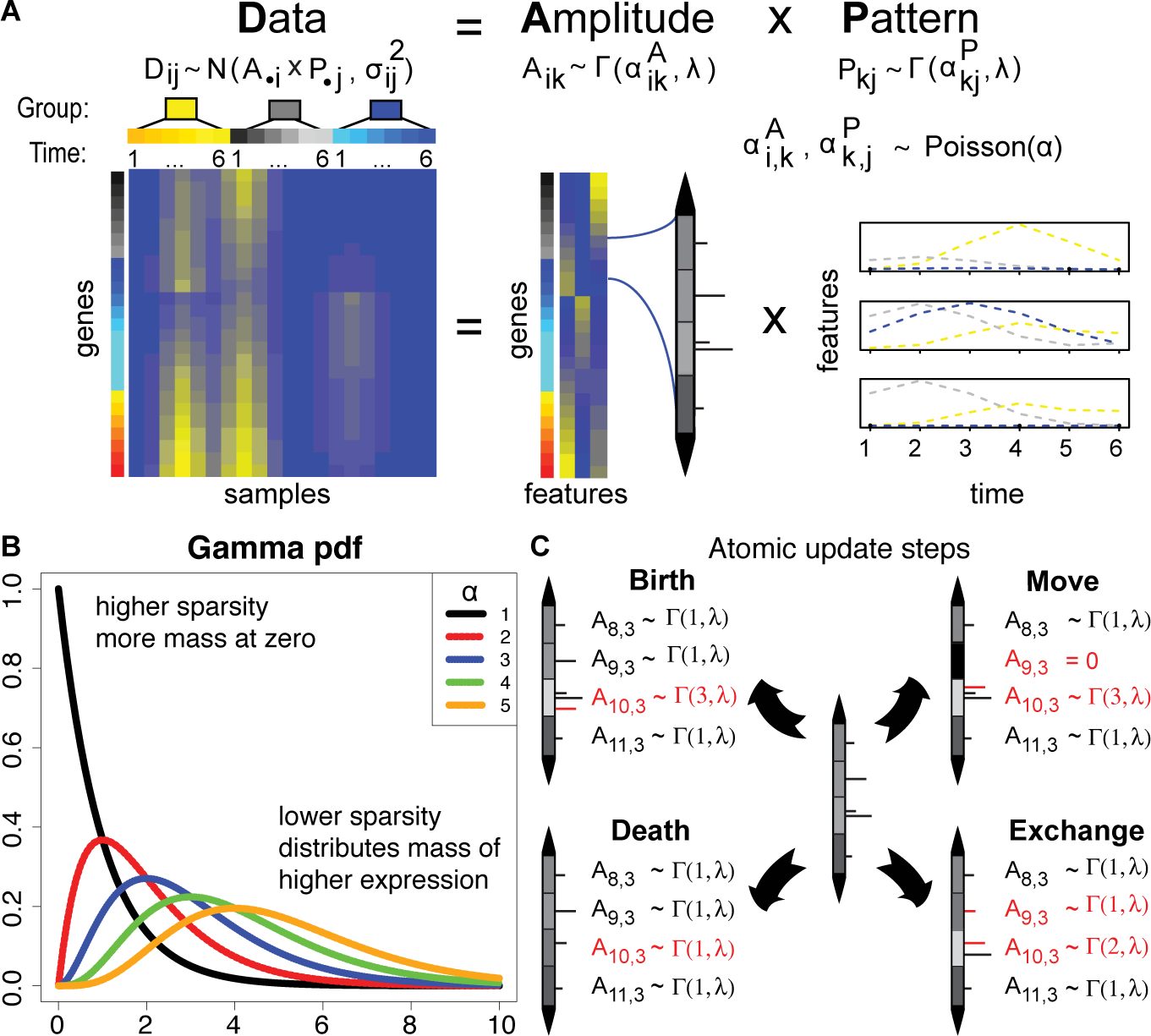
Mathematical core of the scCoGAPS algorithm. (A) scRNA-Seq data yields a data matrix that has each sample as a column and each observed gene expression value as a row. scCoGAPS decomposes the preprocessed data matrix into two related matrices. The rows of the amplitude matrix (**A**)quantify the sources of variation among the genes and the columns of the pattern matrix (**P**) quantify the sources of variation among the cells. The matrix product of **A** and **P** approximates the preprocessed input data matrix. The number of columns of **A** equals the number of rows in **P**, and represents the number of dimensions in the low-dimensional representation of the data. Theoretically, each column in the amplitude matrix and the corresponding row of the pattern matrix represents a distinct source of biological, experimental, or technical variation in each cell. The values in the column of the amplitude matrix then represent the relative weight of each gene and the values in the row of the pattern matrix its relative role in each cell. (B)Adaptive sparsity is achieved by placing a Poisson prior on the shape parameter in the gamma distribution for each matrix element (α_*Ai,j*_,α_*Pi,j*_) and a fixed scale parameter for all matrix elements (λ_*A*_ and λ_*P*_) in **A** and **P**, respectively. In expectation, smaller values of α_*i,j*_ will result in smaller values of corresponding matrix element, and vice versa for larger values which will also have a decreased probability of being zero. (C) The expectation on all atoms are updated at each iteration of the MCMC. During the update, the probability of selecting birth or death is selected based on the Poison prior reinforcing the adaptive sparsity.

In order to ensure that these factors appropriately model biology, additional restrictions to the models used to solve for the values of **A** and **P** can embed further biological assumptions and technical errors specific to each genomics data modality. Bayesian NMF techniques can embed biological and technical structure in the data in prior distributions on the **A** and **P** matrices (Kossenkov et al., 2007; Ochs and Fertig, 2012). To accomplish this for bulk data, we previously developed the Bayesian NMF Coordinated Gene Activity in Pattern Sets (CoGAPS) method (Fertig et al., 2010). CoGAPS uses an atomic prior (Sibisi and Skilling, 1996; Skilling and Sibisi, 1996) to model three biological constraints: non-negativity reflective of pleiotropy, sparsity reflective of parsimony, and smoothness reflective of gene co-regulation and smooth dynamic transitions. The first constraint of non-negativity is common to all NMF approaches, while the latter two are specific to CoGAPS. Other sparse NMF approaches also penalize large or non-zero matrix elements. Yet, the atomic prior in CoGAPS is unique in enforcing a sample- and gene-specific sparsity constraint, which we term “adaptive sparsity”.

In the atomic prior, each element of the **A** and **P** matrices is either zero or follows a gamma distribution (Fig 1A). The adaptive sparsity is achieved by placing a Poisson prior on the discrete shape parameter in the gamma distribution for each matrix element (α_*Ai,j*_,α_*Pi,j*_) and a fixed scale parameter for all matrix elements (λ_*A*_ and λ_*P*_) in **A** and **P**, respectively. Smaller values of α_*i,j*_ result in smaller values of the corresponding matrix elements, and vice versa for larger values. Thus, in some cases the sparsity constraint on values of latent factors will be relaxed in this model, constraining some matrix elements away from zero (Fig 1B). Adaptive sparsity can also model biological structure in the presence of the technical dropouts and true biological zeros in scRNA-Seq. To accommodate the additional sparsity of scRNA-Seq data, λ_*A*_ and λ_*P*_ are set as proportional to the mean of all non-zero values in the data. In contrast, λ_*A*_ and λ_*P*_ for bulk RNA-Seq data are set using the means of the entire data set. The gamma distribution with a discrete shape parameter k is a sum of k exponential distributions, which enables efficient Gibbs sampling with this sparsity constraint in CoGAPS (Supplemental Methods). This formulation also enables modeling smoothness by grouping closely related dimensions near each other. This is achieved by so-called move and exchange steps that shift a single exponential between adjacent matrix elements (Fig 1C). In practice, this step retains the global Poisson prior on shape and the gamma prior on matrix elements while altering the shape parameters between adjacent matrix elements to model smoothness.

### Parallelization and data structures for cross-validation and efficiency: Theory

As a class, Bayesian NMF algorithms such as CoGAPS have substantial computing costs that limit their application to the large datasets generated as tissue atlases with scRNA-Seq data. As we describe in the Supplemental Methods, representing the gamma distribution as a sum of exponentials enables efficient Gibbs sampling. We couple this representation with new data structures for their storage and corresponding calculations that are more efficient than previous versions of CoGAPS and greatly reduce the computational cost for scRNA-Seq analysis (Fig S1A).

We can leverage our hypothesis that latent spaces learned from scRNA-Seq data are reflective of relative gene use in biological processes to enhance the efficiency of Bayesian NMF methods. In this case, distinct subsets of cells sampled from the same condition will have similar factors in a latent space, similar to our previous observation of similar factors across distinct subsets of genes in bulk data (Stein-O’Brien et al., 2017a). The latent space inference with Bayesian NMF can then be run in parallel for distinct subsets of cells in the input scRNA-seq data. We selected the sets of cells in each set based upon the cell-specific composition to enable inference of latent space factors in highly skewed distributions of samples as can occur with rare cell types. As a result, this approach is formally a semi-supervised method in which inference of gene weights in factors are unsupervised. Consensus factors are then created across the sets as described previously for random sets of genes (Stein-O’Brien et al., 2017a). In addition to gaining efficiency, the factors estimated parallel across subsets of cells can also be compared to enable cross-validation of the inferred latent spaces (Fig S1B).

### Transfer learning via dimension reduction using projectR: Theory

Our model that the learned patterns comprising the lower dimensional space describe known and latent factors of the biological system can be used to compare independent, biologically related datasets. This comparison is made by defining a function from the factors in one dataset and an independent, biologically related target dataset into a lower dimensional space that is common to both. Projection is defined as a mapping or transformation of points from one space to another, often a lower-dimensional space. Mathematically, this can be described as a function ϕ(x)=y: R^D^⟶R^d^ s.t for d≤D, x∈R^D^, y∈R^d^. The innovation of projectR is the use of a mapping function defined from the factors in a source data set which enables the transfer of associated cellular phenotypes, annotations, and other metadata to samples in the target dataset (Fig 2).

**Figure 2.**
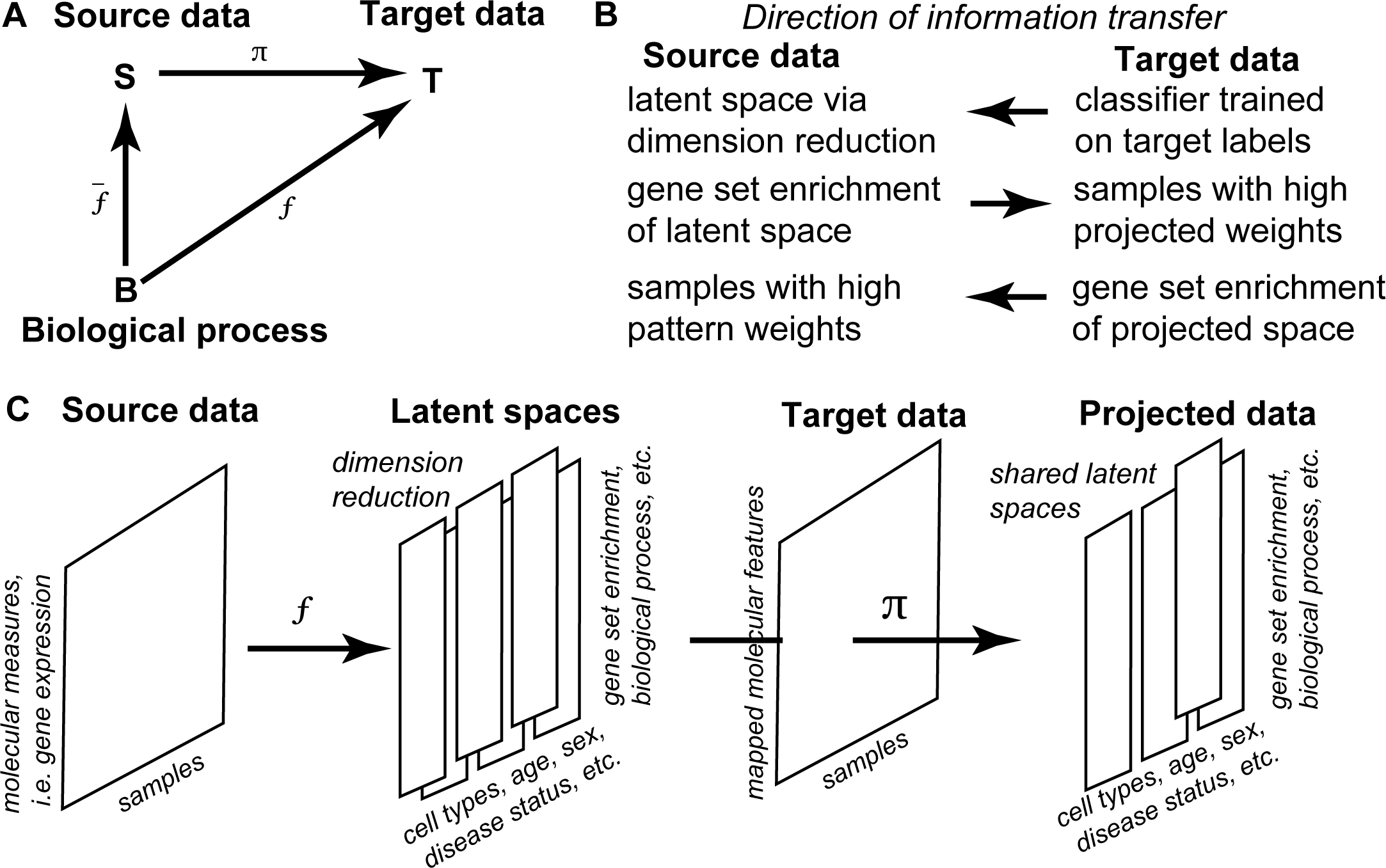
Theoretical core of the projectR algorithm. (A) Graphical representation of projection implemented in projectR showing the relationship between the learned functions, or mappings, and the datasets being operated on. (B) Examples of type and directionality of knowledge transfer enabled via projectR. (C) Diagram of the pipeline used to first learn latent spaces and then project them to transfer learning as describe.

We propose that projection of well-defined latent spaces should capture shared biology across independent datasets. In this study, we perform projection in the column space defined by the amplitude matrix from scCoGAPS (factors representing gene weights), which is accomplished by estimating the patterns P associated with the amplitude matrix by a generalized least-squares fit to the target data (Fertig et al., 2013a) (Methods). Assuming that a given dimension is associated with a specific biological process in the target dataset, the magnitude of the value in this source dataset can indicate its presence within the target dataset. The significance of each projected pattern can be calculated using a Wald test for each sample. Depending on the distribution of the projected sample weights, statistical comparisons between annotated groups can be performed to quantify the presence of these inferred processes in the target data. For example, the mean projected pattern weight between two groups can be compared using standard t-tests or regression-based contrasts. Additionally, classifiers can be built using the projected pattern weights, and the predictive value of each pattern assessed globally. This information transfer enables rapid and highly scalable comparison of very different datasets through the lenses of a projected latent space learned in a source dataset. This analysis can leverage the massive amount of publicly available data and their metadata to annotate the phenotypes in the source data more efficiently. Further, the ability to evaluate whether the processes described by latent spaces are shared, despite significant overall differences in the original high dimensional datasets, can enable hypothesis generation and integrated analyses.

## APPLICATION

### Assessing latent spaces and dimensionality: lessons from bulk RNA-Seq

The developing mammalian retina provides an ideal model system to evaluate the degree to which latent spaces reflect known developmental biology. Features such as discrete cell type signatures, continuous state transitions, signaling pathway usage, developmental age, and sex should each be represented in independent latent spaces. An open question in retinal development is how progenitor cells can generate specific subtypes of neuronal and glial cell types during specific intervals during development—a phenomenon known as progenitor competence (Bassett and Wallace, 2012; Javed and Cayouette, 2017). In an effort to identify genes associated with changes in retinal progenitor cell (RPC) competence, we performed bulk RNA-Seq analysis on replicate populations of FACS-isolated RPCs and post-mitotic cells, using the Chx10:GFP reporter (Rowan and Cepko, 2004), and assessed the fidelity of patterns learned in this bulk analysis across other experimental contexts.

FACS-sorted Chx10:GFP+ RPCs and Chx10:GFP- post-mitotic retinal neurons (Rowan and Cepko, 2004) were collected from the developing mouse retina at three time points, Embryonic day 14 (E14), Embryonic day (E18), Postnatal day 2 (P2), and subjected to standard bulk RNA sequencing (Zibetti et al., 2017). We applied our previous genome wide GWCoGAPS pipeline for bulk RNA-Seq to the normalized FPKM gene expression estimates to identify a latent space consisting of 10 patterns of co-regulated genes (Stein-O’Brien et al., 2017b). Latent space dimensionality can be optimized by maximizing the robustness of patterns between dimensions (Moloshok et al., 2002). Moreover, hierarchies of cell types or subtypes can be resolved by comparing patterns across dimensions (Fertig et al., 2013a). Therefore, we applied GWCoGAPS to the bulk data using a range of dimensionalizations to identify patterns associated with specific biological features or cellular states. Final dimensionality was assessed by comparing factorizations of different dimensions using the ClutrFree (Bidaut and Ochs, 2004) algorithm (Methods). Patterns were strongly correlated (r2>.7) between factorizations at different dimensions, indicating the overall robustness of the factors across dimensions (Fig S1C). For example, a pattern broadly associated with all retinal neurons at a lower dimensionality split into two patterns describing photoreceptors and inner retinal cells at a higher dimensionality as assessed by correlation of cell type specific marker gene expression with individual patterns.

We next evaluated whether patterns identified from bulk RNA-Seq patterns could describe discrete cell type signatures obtained from a comprehensive single cell RNA-Seq dataset conducted across retinal development (Clark et al., 2018). In this study, we isolated 120,804 individual cells from whole mouse retina at 10 developmental time points, ranging from embryonic day 11 (E11) to postnatal day 14 (P14). Single cell RNA-Seq gene expression profiles were obtained using the 10x Genomics Chromium platform (Clark et al., 2018). To relate the data sets, the scRNA-Seq data was projected into the factors learned from the bulk RNA-Seq using projectR (Methods). Using the expert-curated cell type annotations for each single cell, a random forest classifier was trained using projected sample weights as features. Sensitivity and specificity scores were calculated for the relationship between each bulk factor and the annotated cell types detected using scRNA-Seq.

While few patterns had high AUC values for specific cell types, most had moderate values spread across multiple lineages (Fig S1D). One potential explanation for this is that features shared across multiple cells types might dominate the latent spaces found at lower dimensionalization. This finding is consistent with observation that highly expressed unique genes tend to dominate differential expression analysis in bulk RNA-Seq (Ching et al., 2014). An alternative hypothesis is that latent spaces learned in aggregate bulk measures may not cleanly define discrete cell types. As bulk RNA-Seq is inherently an aggregation, testing these hypotheses requires independent measures of each cell. As scRNA-Seq allows for individual measurements of distinct cells, finding similar latent spaces directly from these data would provide strong evidence of their biological vs. technical source. To test this, we next applied scCoGAPS to learn latent spaces directly from the developing mouse retina scRNA-Seq dataset.

### ScCoGAPS identifies signatures of cell types and continuous biological processes in the developing retina

Latent spaces can also be learned directly from scRNA-seq data without projection. Instead of decomposing processes from tissue-level samples, the latent spaces learned across a population of individual cells can identify shared molecular states as well as discrete cell types. Unlike other single cell matrix deconvolution methods that learn factors on a subset of individual cells and then extrapolate to the entire dataset (Macosko et al., 2015), scCoGAPS uses an ensemble-based approach to learn factors in parallel across all of the cells in a dataset, regardless of size. Although not required for this approach, a sampling scheme using a priori curated cell type annotations ensures representation of rare cell types in each set. This sampling enhances pattern discovery from low-abundance cell types across multiple sets. Still, pattern robustness could be assessed by comparing the factors learned across ensembles for biological processes, common cell types not biased by the sampling scheme, and gene set enrichment analysis of the gene weights in the amplitude matrix that are associated with each factor.

ScCoGAPS analysis was performed across the log-transformed, normalized copies per cell from the developing mouse retina single cell RNA-Seq study (Fig 3A)(Clark et al., 2018) and identified 80 consensus patterns (Fig S2). Patterns were learned on a previously described subset of high-variance genes (Clark et al., 2018). Pattern weights from the P matrix were tested for predictive power (AUC) for each cell type annotation. Learned patterns corresponded to both discrete cell types and to continuous state transitions including cycling retinal progenitor populations, a transient neurogenic phase, and intervals of cell type-specific maturation along developmental trajectories (Fig S1F). By associating genes with multiple patterns, scCoGAPS distinguishes gene regulatory modules that are both cell type-specific as well as identified across multiple cell types, thus reflecting shared biological processes (Fig 3B,C). For example, a subset of the shared patterns are active in the early-born retinal precursor cells (RPCs) and another subset in the later-born RPCs (Fig 3D), which were divided into two transcriptionally distinct developmental epochs in our previous study (Clark et al., 2018). Additionally, many shared patterns only account for a small proportion of the cells in later-developing populations, suggesting that these transcriptional programs may be transient or describe features associated with a subset of cells in a given lineage. Comparing pattern weights to human-curated annotations identified several states and conditions which could not be directly inferred from existing annotation alone (Fig 3B, Fig S2), suggesting additional functional relationships exist between cells beyond the annotated relationships described.

**Figure 3.**
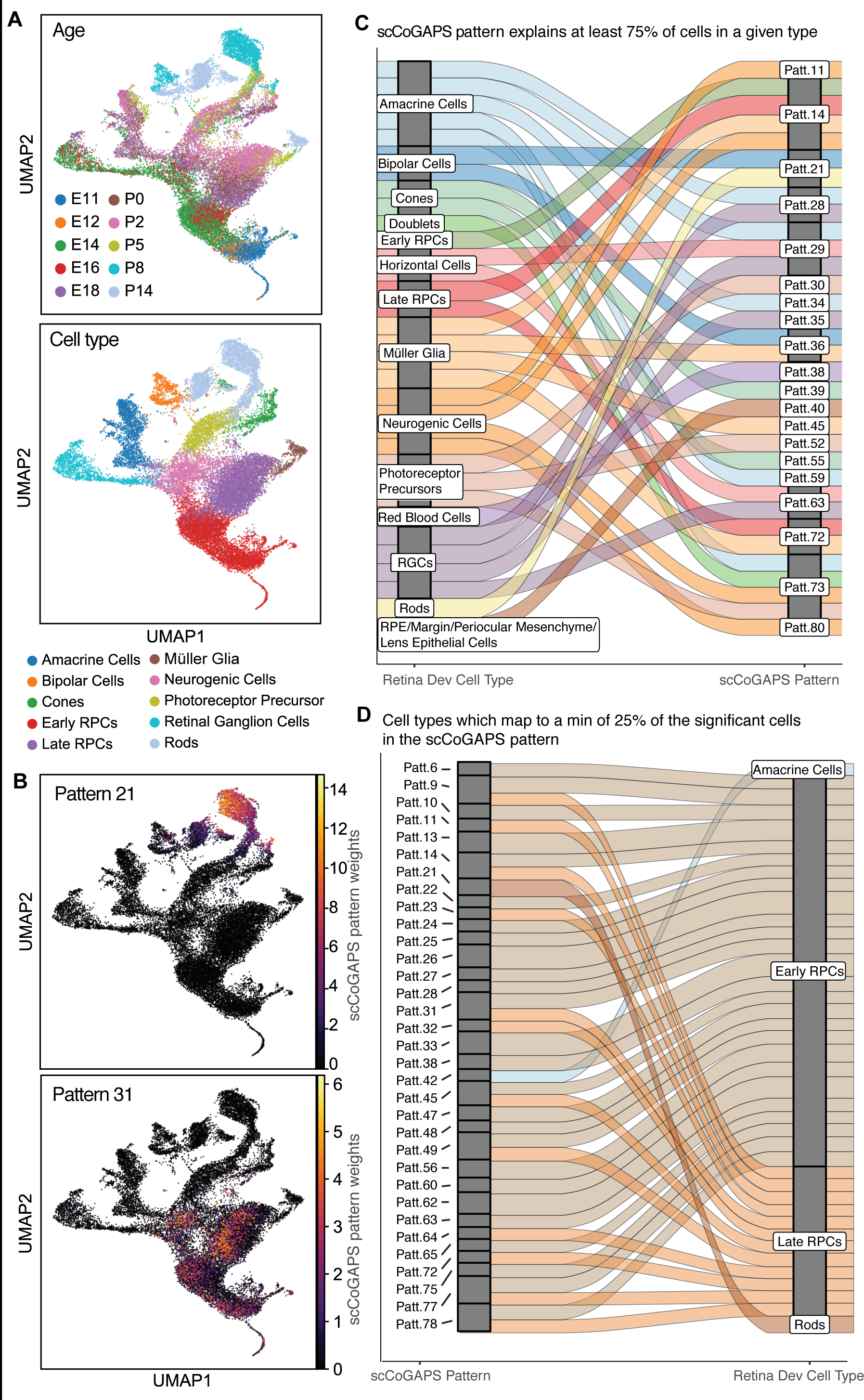
scCoGAPS analysis of time course scRNA-Seq data from developing mouse retina. (A) UMAP of scRNA- seq colored by age (top) and human annotated cell types (bottom). (B) UMAP of retina development colored by scCoGAPS pattern weights illustrate cell type specific (rods, top) and shared (cell cycle, bottom) patterns. (C) Alluvial of cell type specific patterns links manually annotated cell types to scCoGAPS patterns for which at least 75% of cell of a given type have a pattern weight of >0.01. (D) Reversing the alluvial to connect scCoGAPS patterns to cell types for which at least 25% of all cells in a given pattern have a pattern weight of >0.01 demonstrates that patterns shared across multiple cells types and/ or which describe continuous biological processes have strong representation in the progenitor populations. Patterns 21 and 42, which are strongly associated with rods and amacrine cells, respectively, are exceptions to this trend.

To further explore the biological processes captured in each pattern, we analyzed the gene weights (A-matrix and their uncertainty) for each learned pattern. Gene Ontology (GO) enrichment analysis was performed using the CoGAPS gene set test (Fertig et al., 2013b) across all Kyoto Encyclopedia of Genes and Genomes (KEGG) and GO gene sets with <100 genes (Fig S1E, S3, S4). Strong cell type-specific patterns observed included endothelial-specific patterns (9,10 and 56) which are associated with angiogenesis and blood vessel patterning, as well as microglial patterns (5, 6, 24, 25, 27, 57, and 58), which all showed significant enrichment for immune cell activities and processes (p < 1×10-6, Fig S4, Table S4). Patterns specific to neuroretinal-derived cells were also enriched for expected ontologies. Concordant with their selective expression in rods and cone photoreceptors, respectively, patterns 21 and 39 are enriched in phototransduction, visual perception, and photoreceptor cell maintenance, photoreceptor outer segment terms (p < 1×10-8, Fig S4, Table S4). RPC-associated patterns (13, 26, 31, 33, 45, 49, 62, 64, 72, and 78), are enriched for cell cycle regulators and embryonic development as expected (p < 1×10-8, Fig S4, Table S4). Finally, since RGCs are the only neuroretinal cells that extend long projection axons and the only cell to undergo high rates of apoptotic cell death during mouse retinal development (Young, 1984), it is likewise not surprising that the RGC-associated Patterns 15 and 35 are enriched for axon guidance, with Pattern 15 also enriched for negative regulation of apoptosis.

### Transfer learning of retina cell types via projection analysis

To demonstrate the utility of projectR to transfer signatures of cell types and biological processes across datasets, we performed a projection analysis against a separate scRNA-Seq dataset. Specifically, we used our developing retinal dataset as the source data and compared it to a previously published single cell RNA-Seq dataset from P14 mouse retina, established using a different droplet-available time course analysis of human bulk RNA-Seq from whole retinas into our single cell scCoGAPS patterns from mouse retinal development. Homologous genes were used to map the amplitude values across species (Supplemental Methods). Briefly, log2-transformed gene expression values from human retina bulk RNA-Seq data from gestational day 52 to 136 were projected into the 80 mouse developing patterns. Each projected pattern was evaluated for predictive potential for a given human developmental time point with the expectation that the changes in predictive power should reflect the change in pattern utilization over human retinal development. The resulting AUC values for the projected pattern weights revealed a temporal gradient for highly cell type-specific patterns, which reflects both developmental age and relative abundance of each cell type in the bulk sample (Fig 5A). Furthermore, the stereotyped birth order of major retinal cell types (Clark et al., 2018) was faithfully recapitulated in the progression of cell-type-specific pattern in the human time course.

The observed gradient reflects the previously reported three major gene expression epochs of human retina development (Hoshino et al., 2017). The first epoch includes genes with high expression from gestational day (D)52 to D67 and then rapidly downregulated. Patterns associated with early born cell types such as horizontal cells (pattern 1) and RGCs (pattern 15) peaked early (days 57 and 67, respectively) and then declined, reflecting their decreasing relative abundance as later-born cell types are generated. Patterns with amplitude values significantly enriched in RPC-specific processes such as cell cycle regulation (pattern 31) exhibited significant projection in the first epoch (Wald test; BH-correction; q < .01) with AUC values greater than .7 as well. Furthermore, the increased resolution of the patterns derived from scRNA-Seq allowed for a more granular association of corresponding biological processes within the larger epoch. These results indicate that shared continuous features associated with developmental programs in both mouse and human retinal development can be identified via transfer learning with projectR.

Species specific differences were also apparent in this projection analysis. For example, genes that mark mature cone and rod photoreceptors are strongly expressed postnatally in mice (Blackshaw et al., 2001, 2004; O’Brien et al., 2003) but are detected prenatally in humans. Consistent with these previous observations, patterns 39 and 21, which were associated with mouse cones and rod cell types, respectively, exhibit high AUC values during the third epoch of gene expression in our human projection analysis (Fig 5A) (Hoshino et al., 2017). Previous analysis of the bulk RNA-Seq data had demonstrated that differentially expressed genes within the third epoch were enriched for gene ontology terms related to photoreceptors, synaptic connectivity, and neurotransmission (Hoshino et al., 2017). Mouse homologs of the genes annotated with these GO terms were also significantly enriched in the higher amplitude values of our scCoGAPS source patterns 39 and 21 (p <.001) confirming that projectR recovered the species-specific temporal based technique (Macosko et al., 2015), as the target data. The target Drop-Seq single cell dataset was projected into the space of the 80 scCoGAPS patterns from the source 10x-based retinal development time-course data.

**Figure 4.**
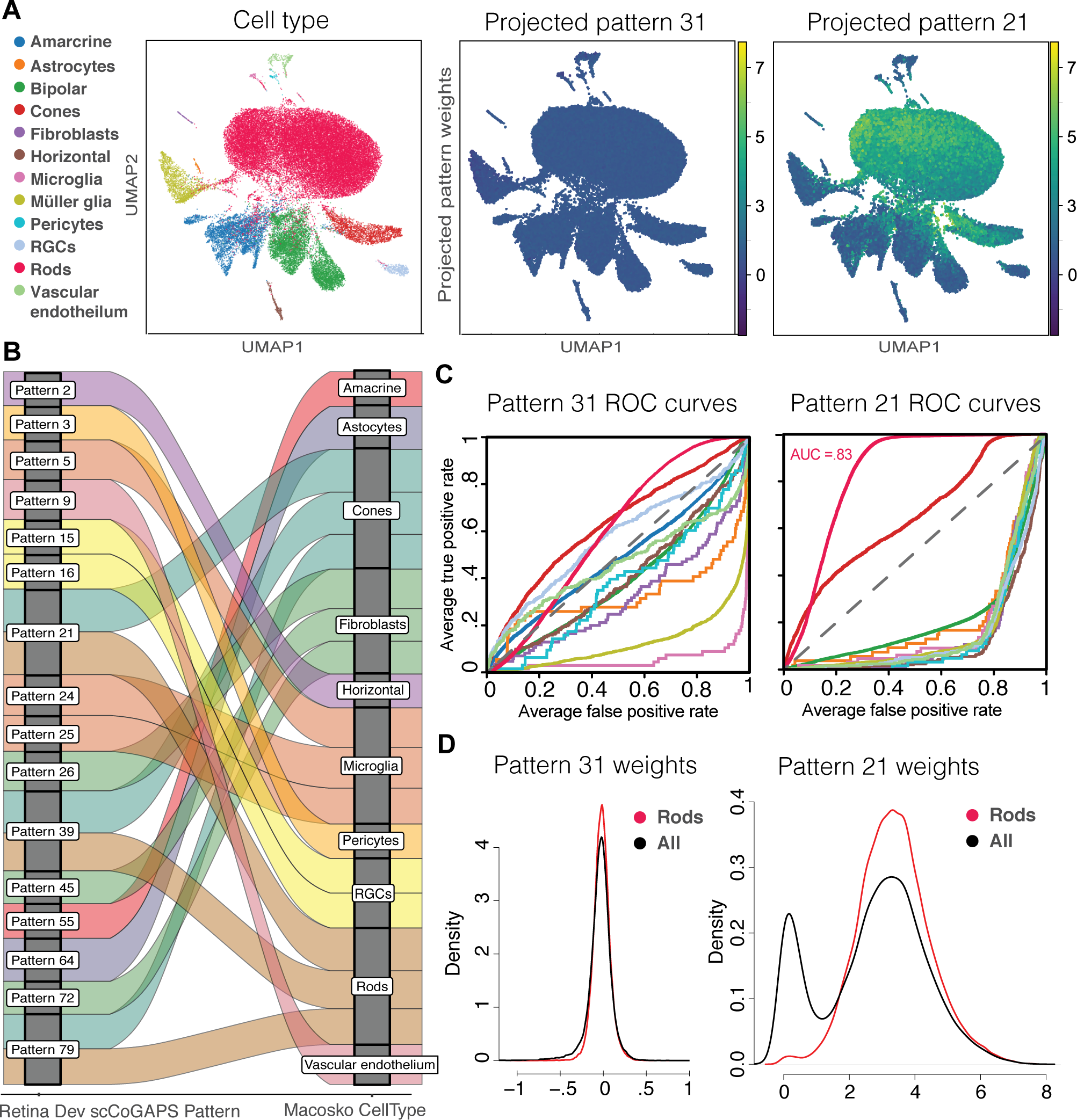
projectR recovers shared cell types in independent murine retina scRNA-Seq data. (A) UMAP of DropSeq data from P14 mouse retina colored by annotated cell type (left), projected pattern weights in Pattern 31 (center), and projected pattern weights in pattern 21 (right). (B) Alluvial plot of projected patterns links previously annotated cell types to scCoGAPS patterns for which at least 75% of cell of a given type have a significant projection (Wald test; BH-correction; q < .01). (C) ROC curves for classifiers built using the projected pattern weights for pattern 21 (right) and projected pattern weights in Pattern 31 (left). Cell types are colored according to the legend in panel A. (D) Density plots of projected pattern weights for all cell types (black) and rods only (red).

**Figure 5.**
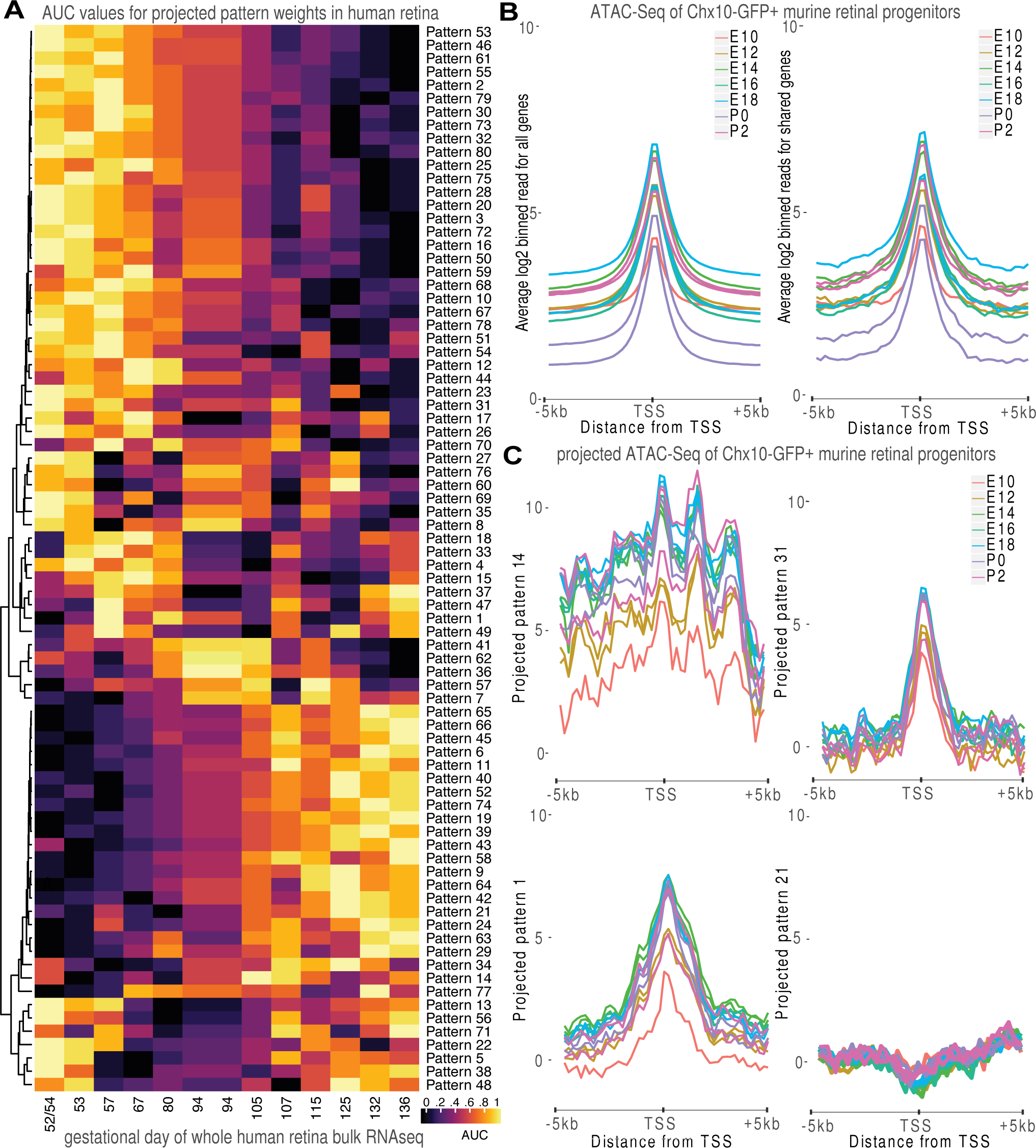
Projection of retina time course data reveals shared temporal dynamics across species and platforms. (A) Heatmap of AUC values for projected pattern weights in developing whole human retina recapitulates previously established gene expression epochs. (B) Average ATAC signal for binned read counts overlapping 200 bp interval extending out 5kb on either side of the transcription start for all genes (left) or the subset of genes from which the scCoGAPS patterns were learned (right). (C) Projection of binned read counts overlapping 200 bp interval extending out 5kb on either side of the transcription start into scCoGAPS patterns 14 (top left), 31 (top right), 1 (bottom left), and 21 (bottom right).

We hypothesized that shared factors would stratify target data consistent with their underlying biological processes, while artifacts or data-specific features would not. AUC values were calculated for each cell type in each pattern using annotations derived from previous analysis (Macosko et al., 2015) (Fig S5A). Consistent with our hypothesis, AUC values confirm that patterns associated with mature cell types present in both the source and target dataset have significant predictive power (AUCs >.7, Wald test; BH-correction; q < .01), while those patterns associated with developmental processes only in the source data did not exhibit significant projections in the more mature (P14) target dataset (AUC <.7, Wald test; BH-correction; q > .01) (Sing et al., 2005). For example, pattern 21, which was strongly associated with rods in the retina development time-course data, selectively marked rod photoreceptors in the P14 retina Drop-Seq data (Fig 4A right panel; AUC = 0.83). Other patterns of mature cell types included pattern 2 (AUC of 0.95 for Horizontal cells), pattern 55 (AUC of 0.91 for Amacrine cells), Pattern 15 and 16 (AUC of .93 and .92, respectively, for RGCs), and pattern 64 (AUC of .99 for Astrocytes) (Fig 4B). In contrast, pattern 31, which was strongly enriched for GO terms associated with cell cycle and progenitor populations failed to yield any significant signal (Fig 4A middle panel).

Using only the significant patterns associated with mature cell types, we are able to resolve true positive cells from background expression in the target dataset using the distribution of projected pattern weights (Fig 4D). Patterns with poor predictive power, such as pattern 3, exhibited weights centered around zero in this projection analysis, while patterns with high predictive potential, such as the rod-specific Pattern 21, exhibit a bimodal distribution (Fig 4D). Cells in the target dataset annotated as rods, however, exhibit a unimodal distribution overlapping with the higher intensity peak of projected pattern weights. The cells contributing to the lower intensity peak therefore have some degree of the pattern 21 rod signature contributing to their transcriptional profile that likely reflects contamination acquired during dissociation and sample preparation These results validate the biological basis of the scCoGAPS patterns for mature cell types and demonstrate the sensitivity and specificity of projectR as a system to transfer annotations based on factors containing shared biological features across datasets.

### ProjectR recovers continuous processes and temporal progression in retina development from disparate data types and across species

We tested whether projection analysis could identify signatures of more continuous biological features across organisms, such as developmental trajectories, rather than just discrete cell types. Specifically, we projected a publicly differences in the use of these patterns through a purely data-driven process of discovery.

To further test the ability of projectR to resolve temporal patterns, we next projected a separate bulk RNA- Seq time course of dissected regions of the human retina from Hoshino et al. (Hoshino et al., 2017). The fovea/macula has been shown to be developmentally ahead of age-matched nasal central and peripheral retina (Hendrickson and Drucker, 1992; Hendrickson et al., 2012; O’Brien et al., 2003), and enriched for both cone photoreceptors and retinal ganglion cells (Curcio and Allen, 1990). In the original data, a differential gene expression analysis of macula vs periphery was underpowered to detect significantly differentially expressed genes at each time point. However, using the projected values for each sample, we could readily calculate differential pattern usage. Significant differential pattern usage (Wald test; BH-correction across patterns; q<.01) was observed between the fovea/macula and peripheral retina at days 73 and 132. In this projection analysis, we observed that the fovea/macula is enriched in patterns specific to mature neurons, particularly retinal ganglion cells and cones (Patterns 1, 15, 39, 52) and depleted in patterns specific to retinal progenitor cells (Patterns 26, 31, 78) or immature neural precursor cells (patterns 17, 73) relative to the age-matched peripheral retina (Fig S5B). These results demonstrate the utility of projectR in recovering spatiotemporally regulated differences within tissue/organ development.

Projection analysis can also determine pattern usage across a variety of different cellular measurement types. To illustrate this, we determined whether patterns learned from scRNA-Seq analysis of the developing mouse retina could be used to identify distinct chromatin accessibility profiles within a mouse retinal ATAC-Seq time-series obtained from FACS-isolated Chx10:GFP-positive RPCs (Rowan and Cepko, 2004) collected at two day intervals between E10.5 and P2 (Fig 5B-C; Fig S6). Since ATAC-Seq data profiles chromatin accessibility, rather than gene expression per se, projection analysis allowed identification of patterns associated with genes that are primed for transcriptional activation. For each gene, ATAC-Seq reads were quantified in 200 bp bins −5Kb to +5Kb around each canonical TSS for each time point sampled (Methods). As expected, the naïve signal shows global enrichment over TSSs owing to the increased accessibility at TSS of actively transcribed genes (Buenrostro et al., 2013) (Fig 5B). Overall signal intensity was highly variable, with biological replicates from the same time point demonstrating a strong batch effect. These effects persisted when the ATAC-Seq data were subset to the same high-variance genes used to define the scCoGAPS patterns (Fig 5B right). In order to test the ability of projectR to overcome these effects, no batch correction or further data normalization was performed.

Despite the consistent profile of the observed mean enrichment of ATAC-Seq signal at the TSS across sample, projection of the ATAC-Seq into the scCoGAPS patterns revealed several classes of chromatin accessibility patterns. Indeed, different accessibility ‘shapes’ emerged that were previously contributing to the global average, but lost in aggregate. Furthermore, the shape of the accessible peak and ranking of samples was relatively distinct across the different patterns, indicating that projection analysis can recover discrete signatures of accessibility associated with factors learned from gene expression profiles, independent of technical noise. Together, these results suggest that learned accessibility signatures are associated with specific biological processes at distinct developmental timepoints in the developing mouse retina. Specifically, patterns that reflected absent biological processes (such as mature cell types not sampled in the ATAC-Seq) demonstrate no significantly appreciable signal in the projection analysis, while shared processes are apparent in both the scRNA- Seq and the ATAC-Seq data. For those projected patterns with significant ATAC-Seq signal, replicates displayed significantly tighter concordance, and the amplitudes of the projected accessibility signatures appropriately reflected temporal progressions.

Broad domains of open chromatin on either side of the transcriptional start site—a hallmark of strongly transcribed genes—were observed exclusively in patterns associated with proliferating RPCs (e.g. patterns 14,45,72,78), consistent with the ATAC-Seq sampling of this population. Sharp peaks of open chromatin centered on the TSS corresponded to RPC-specific patterns associated with actively transcribed genes (e.g. patterns 4,31,64), as well as a subset of patterns associated with maturing retinal subtypes, including cones, RGCs and ACs (e.g. patterns 1,2,15,39), and immature rod photoreceptors (pattern 79). Finally, TSS signatures of closed chromatin are associated with patterns specific to cells that are not derived from RPCs, such as microglia (5,24) and erythrocytes (28), as well as with the mature rod photoreceptor-specific Pattern 21. These data indicate that promoter regions associated with genes specific to RPC- derived cell types exist in an open and poised state in RPCs, with the notable exception of genes specific to mature rods. Taken together these results reinforce a developmental analysis of chromatin conformation obtained from whole mouse retina (Aldiri et al., 2017) and demonstrate the utility of projectR to recover meaningful biological signal across disparate data types using continuous weighting for each factor.

### projectR enables factor comparison across model systems: from the developing retina to the developing brain

The retina is often used as model system for neural development given its relative accessibility compared to the rest of the CNS and precise organization. In particular, both retinal neurogenesis and corticogenesis share a stereotyped birth order of different lineages from a single progenitor population (Kohwi and Doe, 2013; Miller and Gauthier, 2007). To test the ability of projectR to identify conserved pattern usage across tissues and model systems, we projected our retinal scRNA-Seq patterns into two datasets derived from developing human cortex (Nowakowski et al., 2017) (Zhong et al., 2018), and an additional dataset of the developing mouse midbrain (La Manno et al., 2016) (Fig 6). The projection of these 80 patterns across all cells in each of the datasets completed in 165.6, 56.0, and 3.0 seconds on a high performance computing (HPC) cluster node with a 2.5 GHz AMD Opteron Processor 6380, and 40Gb of RAM. Consistent with a significant degree of conserved developmental processes and tissue composition between retina and select other CNS regions, we identified 87.5% (70/80), 76.3% (61/80), and 98.8% (79/80) of patterns with significant projection (q≤0.01; Wald test) in at least one cell in each of these comparable model systems (Fig 6 & S8), suggesting that many of the processes described by these patterns are reused in other CNS regions.

**Figure 6.**
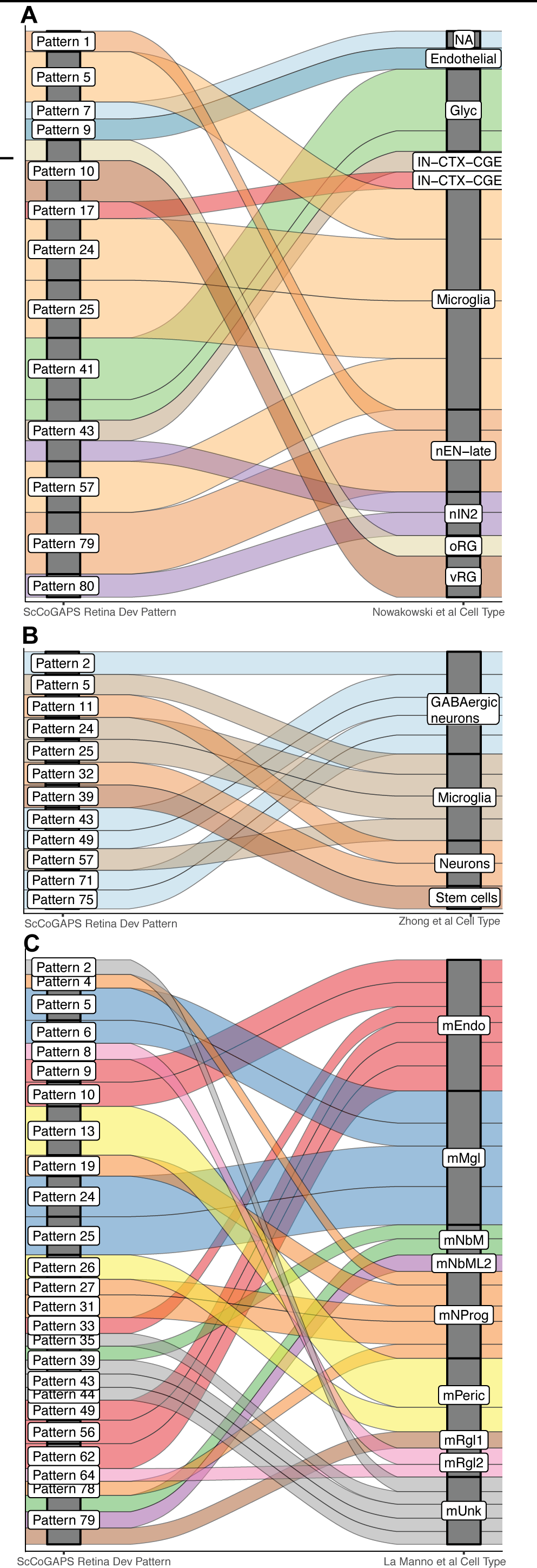
Developing brain scRNA-Seq projected in scCoGAPS patterns of retina development. Alluvial plots connecting scCoGAPS patterns to cell types for which at least 25% of all cells are significant (Wald test; BH-correction; q < .01) in a given a projected scRNAseq of human cortical development from (A) Nowakowski, et al. and (B) Zhong, et al. as well as projected scRNAseq of mouse mid-brain development from (C) La Manno et al.

For the human cortical data, patterns 5, 20, 28, 29, 31, 40, 53, 64, and 65 captured 75% of published annotated cell types (Fig S7A). Consistent with its derivation as a progenitor-associated pattern in the developing retina and GO enrichment for cell cycle, pattern 31 demonstrated significant (AUC >0.7; q≤0.01; Wald Test; BH-corrected) projection to basal intermediate progenitor cells (IPCs), IPC-derived neuronal precursors of the medial ganglionic eminence (MGE), and dividing radial glia in the cortex (Fig S8A). Notably, in cortical data from Nowakowski, et al., we observed that pattern 43, which is specific to inhibitory amacrine cells in retina, is also associated with interneurons (Fig 6A & S8A). Newborn excitatory pyramidal neurons are enriched for genes found in both the photoreceptor precursor-enriched pattern 79 (*Unc119, Meis2, Cdc43ep3*), as well as the amacrine and horizontal cell-enriched pattern 1 (*Nrxn3, Kdm5b, Dusp1*). Additionally, we are able to classify the unannotated cells (NA) as neurons using projection of pattern 7 which is enriched for mature neuronal markers (*Nnat, Tubb2b, Nefl*). In data from Zhong et al., where progenitors and precursors of GABAergic interneurons are annotated as a single class, these cells were significantly associated with patterns specific to GABAergic horizontal and amacrine cells (2,43) and RPCs (49,71) (Fig 6B). In the mouse midbrain, neural progenitor cells were enriched for retinal progenitor-specific patterns 4, 31, and 78, consistent with their shared roles in these two tissues (Fig 6C). Notably, Glyc cells in human cortex and mUnk cells in mouse midbrain—neither of which could be confidently classified in the original studies—are both enriched for patterns and genes (*Tubb2b, Sox4, Mapt, Onecut2*) specific to immature amacrine, horizontal and/ or RGC cells, indicating that these both represent as yet unidentified neuronal precursor subtypes (Fig 6C). These associations further demonstrate that projection analysis can be used to identify and annotate comparable cell types and shared biological processes across disparate model systems, and that information transfer faithfully recovers these associations across species (Fig S8).

Patterns 5, 6, 24, 25, and 57 are each associated with microglial cells in the original source dataset. Interestingly, we observed several significant differences in the projections of these patterns into microglia from different CNS regions, as well as across species. Patterns 5, 24, and 25 were consistently associated with microglia in all three brain region projections(Fig 6A-C). However, pattern 57, was significantly (q<0.01; Wald test; BH-corrected) associated with microglia in both human cortical projections, but not in microglia from the mouse midbrain (Fig 6A-B, Fig S7A-B), suggesting a potential difference in microglia signatures derived from different CNS regions. This pattern projection is driven in part by the Cathepsin family member genes *Ctsb* and *Ctsd*, as well as *Cd9*, each of which has been previously shown to be upregulated in a subclass of cortical microglia (Keren-Shaul et al., 2017). These results suggest that pattern 57 may be specifically associated with the cortically-enriched microglia type II, and highlighting a region-specific property of microglia detected via projection analysis. Additionally, no significant projections for pattern 6 were identified in either human CNS dataset (Fig 6C, Fig S7C); 0/68 (0%) annotated microglia in Zhong et al. and 0/77 (0%) microglia in Nowakowski et al. demonstrated a projection value with q<0.01. This is in contrast to 76/77 (98.7%) microglia in the human cortical development study with significant (q≤0.01; Wald test; BH-corrected) projections into the other microglial pattern 5. Thus, using projectR we are able to discriminate region- and species-specific differences in the transcriptional signatures of discrete cell types.

### Shared latent spaces identify novel cell type associations across a single cell atlas of adult mouse tissues

Given that latent spaces may reflect the signatures of biological processes in the conditions in which they are learned, we next asked whether we could identify significant use of these processes in other diverse cellular contexts from an atlas of adult mouse tissue single cell RNA-Seq. The Tabula Muris dataset is a publicly available collection of 70,118 single cell gene expression profiles from 12 mouse tissues (Wyss-Coray et al., 2018) collected using the 10x Genomics Chromium platform (Fig 7A). Using projectR, we projected the Tabula Muris dataset into the latent spaces of the scCoGAPS patterns. This analysis completed using all cells in 107 seconds on a HPC cluster node with a 2.5 GHz AMD Opteron Processor 6380 and 40Gb of RAM. Consistent with our hypothesis that biologically meaningful latent spaces will be shared across diverse cell types, 83.8% (67/80) of the patterns demonstrated significant projection (q≤0.0001; Wald test) in at least one cell, and significant projections were identified in each of the 12 tissues in the Tabula Muris dataset.

**Figure 7.**
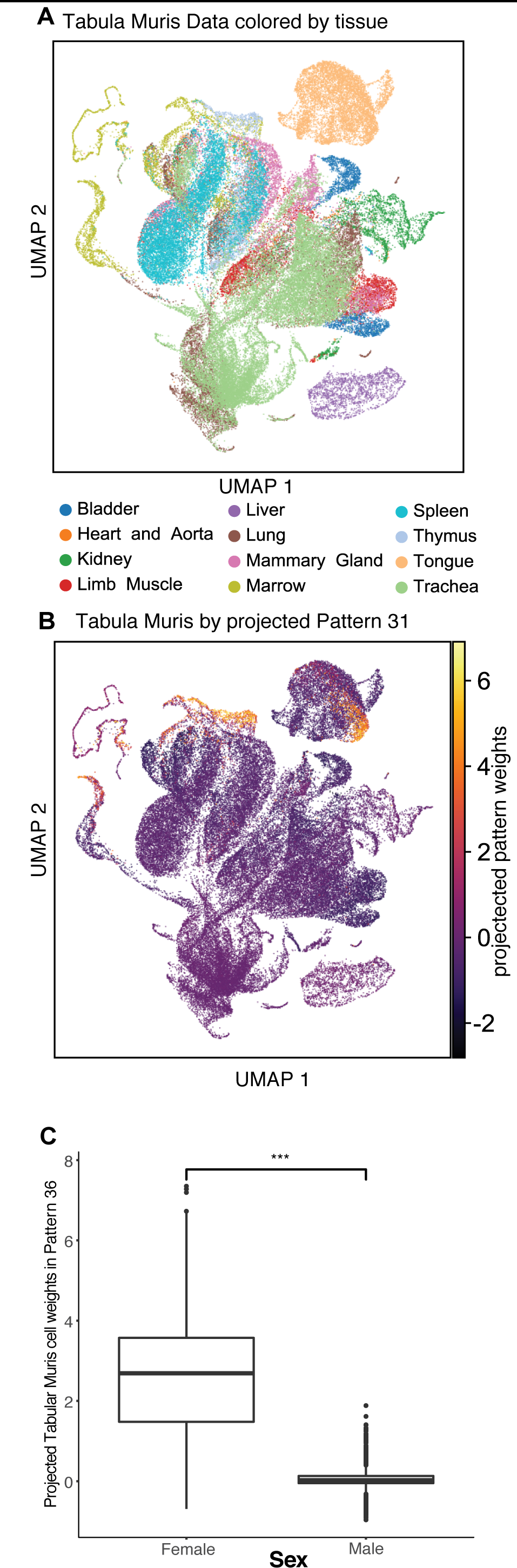
Projection of retinal scCoGAPS patterns into mouse non-neuronal cell dataset. (A) UMAP of scRNA-Seq data from the Tabular Muris collection of mouse tissues colored by tissue and (B) projected pattern weights in pattern 31. (C) Boxplot of projected Pattern 36 weights stratified by sex demonstrates statistically significant difference corroborating association with genes involved in X-inactivation.

Using only patterns learned in the developing retina, we were able to identify, characterize, and annotate a variety of cellular features across this broad mouse cell atlas including several obvious cellular phenotypes as well as a few less readily-apparent biological features and cellular subpopulations. For example, many progenitor-associated patterns project into adult tissues with high levels of cell turnover, and specifically within subsets of the cells that are actively proliferating (Fig 7B). Consistent with previous projections and GO enrichment for cell cycle, Pattern 31 is highly predictive of actively mitotic cells, and can even be used as a proliferative index via projection (AUC >.7) in proliferative tissues within the Tabula Muris dataset such as marrow, thymus and tongue epithelium (Fig 7B).

As previously described (Clark et al., 2018), we identified pattern 36 as specifically associated with sex in our developing retinal source dataset. This association was further confirmed by defining biomarkers for each factor, computed using the PatternMarker statistic, which finds the set of genes that are uniquely associated with each factor (Methods) (Stein-O’Brien et al., 2017b) (Supplemental File 3). The sole PatternMarker for pattern 36 was *Xist*. Consistent with this expectation, the projection of the Tabula Muris data set into pattern 36 almost perfectly segregated male and female samples (Fig 7C, p-value < 2.2e-16, two way t-test). While females displayed a range of weights, potentially corresponding to variable levels of X-chromosome inactivation and/or Xist transcript expression, males had uniformly insignificant projected pattern weights. We note that the original pattern 36 in our source data had high weights in a large proportion of cells, independent of their age, cell type, or technical annotation. A three-dimensional UMAP representation with each cell colored by its pattern 36 weight revealed an apparent uniform asymmetry of the distribution of transcriptional profiles of the cells using this pattern (Supplemental Video 1). While reflecting basic biology, the projection of pattern 36 between these two datasets provides an example of how annotations from one dataset could be used to annotate patterns in a gene-independent manner.

Patterns specific to retinal neurons were detected in a number of peripheral tissues (Fig. S9A). In the trachea, *Mgp*-positive goblet cells expressed genes associated with the neuronal cytoskeleton and neurotransmission (Gap43, Sncg, Chgb, Tac1). In the tongue, Krt6a/Krt16- positive epithelial cells of both the filiform papillae (pattern 37) and Krt14-positive cells of the basal layer (pattern 41) selectively expressed genes associated with the neuronal cytoskeleton. In the lung, a small number of cells expressed pattern markers associated with amacrine/horizontal cell-enriched Patterns 16 and 17 (Scg5, Tmsb10, Malat1, H3f3a) (Fig. S9A). This lung subpopulation expressed Ins1 and Ins2, and may thus correspond to a previously uncharacterised subset of pulmonary neuroendocrine cells (Fig. S9B-D). In each of these cases, none of the most highly selective marker genes of these cells types (*Mgp, Krt6a/14/16, Ins1/2*) were themselves expressed in retina, but rather the projected patterns identified more complex similarities in gene expression between these peripheral cell types and retinal cells. These findings illustrate the power of this approach to identify biological processes and cellular features shared between otherwise transcriptionally dissimilar cell types.

## DISCUSSION

Here, we demonstrate that a reduced set of continuous factors is sufficient to accurately describe cellular identity, state, and phenotype not apparent from marker genes alone. Moreover, these dimensions, learned from a source dataset, can be used to rapidly learn biologically meaningful relationships across diverse datasets including different assay technologies, cellular measurements, and species. We present two new algorithms based on this theory. The first, scCoGAPS, identifies factors using a Bayesian non-negative matrix factorization approach that can be parallelized across all cells in a given study and appropriately adapts to the sparsity of scRNA-Seq data. The parallelization allows for a computationally tractable factorization of large datasets, such as those proposed by the Human Cell Atlas Project (Rozenblatt-Rosen et al., 2017). Furthermore, parallelization across cells allows for the independent discovery of patterns across sets of cells/ samples that can be used to assess pattern robustness, establishing confidence in the learned factors. Application of scCoGAPS to time course scRNA-Seq analysis of mouse retina development identified gene expression signatures of discrete cell types, as well as shared gene regulatory networks governing neuronal development.

The second algorithm described here is a regression-based tool, projectR, that facilitates rapid and scalable transfer learning of latent spaces across datasets. Using projectR, very different datasets, such as mouse retina and human cortex, can be compared with respect to specific latent spaces to evaluate whether the processes described by each space may be shared despite significant overall differences in the original high-dimensional datasets. In contrast, existing tools for exploratory analysis of shared cellular phenotypes across single cells (Kiselev et al., 2018) rely on consensus clustering using select discriminating marker genes. By mapping target data into a basis set defined by the source data, projectR allows for the direct evaluation of what is shared between, versus what is unique to, the source and target datasets. Additionally, projectR is able to overcome confounding factors from technical variation such as batch effects. This is especially significant for integrating across molecular features where differences in data dimensionality and distribution currently limit implementation of techniques that enforce alignment to a mutually learned basis set (Butler and Satija, 2017; Wang et al., 2015).

The exponential growth and rapid adoption of large-scale, high-throughput biological assays has generated massive amounts of data. Single cell analyses have added to the complexity and scalability required to analyze these data, as individual experiments can now analyze millions or more individual samples. Latent space analyses have already demonstrated useful applications in single cell analyses for identifying and correcting for technical errors associated with mRNA dropout (Eraslan et al., 2018) or analysis of cell-cell variation (Loos et al., 2018). A key remaining challenge is comparing biologically meaningful molecular features across data sets. Currently, the often discordant and inharmonious nature of biologically distinct datasets, and the significant impact of technical variation, both challenge the ability to make meaningful interpretations from direct comparisons of samples of interest (Tung et al., 2017). Our approach extends these concepts and enables the comparison of these factors across a variety of experimental paradigms and cellular contexts.

ScCoGAPS and projectR enable rapid, sensitive, and scalable transfer of the knowledge contained within scRNA-Seq data through the learning of, and subsequent projection into, biologically meaningful latent spaces. Here, we demonstrate the sensitivity of scCoGAPS pattern discovery and projectR projection analysis to recover shared features and annotations across a variety of data types and experimental conditions. Our approach enabled *de novo* annotation and correction of existing cell type annotations in a target retinal scRNA-Seq study. We have demonstrated the cross-platform and cross-species sensitivity of this approach to identify paralogous cell types in the retina and other CNS tissues, and identify meaningful biological similarities in markedly different cell types in a mouse cell atlas. This transfer learning approach has a wide range of significant applications in the current high-throughput biological space including cell type inference, comparison of factors across distinct cell types or condition (e.g. disease vs normal), identification of unanticipated features in a given biological context, and cross-model and cross-assay integrative analyses. The application of scCoGAPS and projectR allows for exploratory analysis of high-dimensional biological data through the lenses of individual biological processes and enables a shift in how we compare and identify cells beyond reliance on specific marker genes or ensemble molecular identity.

## METHODS

### Animals

Mice were housed in a climate-controlled pathogen free facility, on a 14 hour-10 hour light/dark cycle (07:00 lights on-19:00 lights off). All experimental procedures were preapproved by the Institutional Animal Care and Use Committee of the Johns Hopkins University School of Medicine.

### Bulk RNA-Seq of the developing mouse retina

At select developmental time points, cells were collected from biological replicates of FACS-sorted Chx10-Cre:GFP+ mouse retinas as previously described (Rowan and Cepko, 2004). RNA was isolated using the RNAeasy Mini kit (Qiagen) with on-column DNase treatment. Isolated total RNA was assessed for integrity on the Bioanalyzer 2100 system, and we required a minimum RNA integrity number of 7. RNA- Seq libraries were created using the Illumina TruSeq kit (Illumina), quantified via PicoGreen assay and fragment size distribution was determined using the Bioanalyzer 2100. Libraries were barcoded, pooled, and run on a HiSeq2500 instrument. 75-100bp paired-end reads were mapped to the mouse reference genome (mm10) using Hisat2 (Kim et al., 2015, 2016) to an average sequencing depth of 30.0 million aligned reads per sample. Gene expression estimates for the reference transcriptome (Gencode vM5) and differential testing were performed using Cuffdiff2 (Trapnell et al., 2012) with default parameters. Data are in process at GEO.

### Single-cell RNA-Seq of the developing mouse retina

The developmental time-series of scRNA-seq from mouse retina was performed as described in (Clark et al., 2018). UMAP representations were found on neighbors calculated from the first 32 PCs using scanpy version 1.1 following data processing using the zheng17 recipe in python 3.3.

### ATAC-Seq of the developing mouse retina

Chromatin derived from flow-sorted Chx10:Cre-GFP- positive (Rowan and Cepko, 2004) retinal fractions was processed as previously described (Zibetti et al., 2017). Briefly, chromatin was extracted and processed for Tn5 mediated tagmentation and adapter incorporation, according to the Manufacturer’s protocol (Nextera DNA sample preparation kit, Illumina) at 37°C for 30 min. Reduced-cycle amplification was carried out in presence of compatibly-indexed sequencing adapters. Libraries were quantified using the PicoGreen assay and fragment size distribution was determined using the Bioanalyzer 2100. Up to 4 samples per lane were pooled and run on a HiSeq2500 Illumina sequencer to produce 50 bp paired ends for each sample.

Bowtie2 (version 2.3.2) was used for ATAC-Seq reads alignment to the mouse genome (mm10) (Langmead and Salzberg, 2012). Duplicate reads were removed using Picard tools (version 2.10.7)(Wysoker et al., 2013). Improper mapped reads were removed using samtools (version 1.5). (Li et al., 2009). Read counts for each gene were retrieved using featureCounts program (version 1.5.3). (Liao et al., 2014). Read counts overlapping 200 bp interval extending out 5kb on either side of the transcription start site were generated with custom scripts using bedtools (version 2.26.0)(Quinlan and Hall, 2010). Data are in process at GEO.

### Target public domain datasets

All data was downloaded from GEO with the exception of the Tabular Muris data which was downloaded from https://github.com/czbiohub/tabula-muris and the developing human cortex time course from (Nowakowski et al., 2017) which was downloaded from https://cells.ucsc.edu/?ds=cortex-dev. Accession numbers in order of appearance in the manuscript are GSE63472 (Macosko et al., 2015), GSE104827 (Hoshino et al., 2017), GSE104276 (Zhong et al., 2018), and GSE76381 (La Manno et al., 2016).

### Pattern matching for consensus gene signatures

Hierarchical clustering was done on gene weights from all sets and the resulting dendrogram is cut so that the number of branches is equal to the original number of latent spaces. Each branch then contains the columns(s) of **A** across all of the sets that are most related to each other. Well-dimensionalized data will produce robust patterns such that each branch will contain a single contribution from each of the randomly generated sets. As the additional sparsity can cause large clusters driven predominantly by zeros, the minimum and maximum number of patterns contributing to given branch can be specified with defaults of .5 and 1.5 the number of gene sets, respectively. Branches failing to meet the lower bound are dropped, while those exceeding the upper bound are subjected to additional rounds of hierarchical clustering. Additionally, the minimal correlation to the cluster mean for each patterns within a given branch was specified to be 0.7. Consensus signatures were then constructed for each branch by taking a weighted average of the gene signatures for that branch which pass all the criteria. To ease across pattern comparison, the resulting consensus signatures were scaled to have maxima of one. Pattern weights for all the cells were then learned in parallel from these signatures to ensure reciprocity across all of the sets.

### Gene set analysis of scCoGAPS patterns

Z-scores of gene weights were computed for each pattern in each ensemble by dividing the mean of the **A** matrix estimated across the chain by its standard deviation as previously described (Fertig et al., 2010; Ochs et al., 2009). The resulting matrix of Z-scores is averaged for sets of patterns determined to match in the the ensemble as described above. A Wilcoxon gene set test with the R/ Bioconductor LIMMA package version 3.36.2 (Ritchie et al., 2015) is performed for mouse KEGG and GO sets from the R/Bioconductor packages org.Mm.eg.db version 3.4.0, KEGG.db version 3.2.3, and GO.db version 3.4.0. Gene sets with more than 5 genes and fewer than 100 genes are retained for analysis. P-values for the gene set test are FDR adjusted with Benjamini Hotchberg and available as Table S4.

### projectR analysis

The R package projectR version 0.99.2 (available from https://github.com/genesofeve/projectR) was used to project the scCoGAPS consensus scCoGAPS patterns of the **A** matrix into each of the target datasets. These projection are achieved by solving the factorization D = AP + ε using the least-squares fit to the new data as implemented via a wrapper for the lmFit function in the LIMMA package 3.30.13 (Ritchie et al., 2015). By using the **A** weights as the design matrix for multiple linear regression the estimated coefficients provide **P** matrix values for the new data. These **P**s score the new samples using a gene-wise weighting, provided by the **A**s, for each new pattern. The ranking of the new samples within each pattern are then indicative of the relative strength of their biological features associated with the original pattern. A Wald test to calculate the significance of these coefficients is calculated using the pdf of the negative absolute value of the coefficients scaled by their standard deviation. Code to reproduce these analyses is available at https://github.com/genesofeve/retina_dev_methods.

## ACKNOWLEDGEMENTS

This work was supported by grants from the NIH (R01EY020560 and U01EY027267 to SB, F32EY024201 and K99EY027844 to BSC, R01CA177669, U01CA196390, and U01CA212007 to EJF), the Chan-Zuckerberg Initiative DAF (2018-182718 for QH, 2018-183445 to LAG, and 2018-183444 to EJF), an advised fund of Silicon Valley Community Foundation, the Johns Hopkins University Catalyst (EF & LAG) and Discovery awards (EJF), and the Johns Hopkins University School of Medicine Synergy Award (SB, LAG, & EJF). The authors would like to thank C.A. Berlinicke and D.J. Zack for assistance with FACS analysis, J. Taroni for discussions on transfer learning and low dimensional representations, A. Wolf and F. Theis from the Helmholtz Center, Munich, Germany for productive discussions and introductory scanpy code, the Johns Hopkins Genetic Resources Core Facility for use of the 10x Genomics Single Cell system, and the Hopkins Microarray and Deep Sequencing Core for assistance with sequencing; the CZI Jamboree, K. Korthauer, and A. V. Favorov for invaluable collaborations and discussions; and A. Battle, V. Vasan, and J. Bader for comments on the manuscript.

## AUTHOR CONTRIBUTIONS

GSO’B, BSC, SB, LAG and EJF conceived and directed the study. BSC generated scRNA-Seq data. GSO’B, BSC, LAG and EJF analyzed scRNA-Seq data, with LAG and EJF as senior bioinformaticians. CZ and BSC generated the bulk RNA-Seq, and CZ generated the ATAC-Seq data. GSO’B, TS, LAG and EJF developed scCoGAPS. GSO’B, RS, CC, LAG, and EJF contributed to the development of projectR. QH and GSO’B developed the random forest classifier for the projections of bulk GWCoGAPS patterns. RS and GSO’B developed the general classifier for projected pattern weights included in the projectR package. SL, CZ, JQ, and GSO’B analyzed the ATAC-Seq data. GSO’B, BSC, SB, LAG, and EJF wrote the paper, with input from all co-authors.

## SUPPLEMENTAL FIGURE LEGENDS

**Figure S1. RNA-Seq from Chx10:GFP-positive retinal progenitor cells.** (A) Benchmarking of computational infrastructure demonstrates 30x speed up of version 3.7 vs. 3.6 (B) Hierarchical clustering of individual parallelized scCoGAPS solutions. Colors demark sets which will contribute to the same consensus amplitude in the final result. (C) Dimensionality assessment of GWCoGAPS analysis using the Clutrfree algorithm reveals hierarchical relationship of higher dimensional solutions to lower dimensional ones. (D) Heatmap of AUC valued for random forest classifier based on scRNA-Seq data projected into bulk patterns. (E) Distribution of scCoGAPS amplitude values for patterns 21 (top) and 31 (bottom). Genes used for GO enrichment for GO:0007049 cell cycle (blue) and GO:0007602 phototransduction, visible light (orange) are highlighted. (F) Heatmap of AUC values of curated cell types within the mouse retinal development scRNA-seq data for scCoGAPS patterns from mouse retina development

**Figure S2. 80 scCoGAPS patterns from 10x RNA-Seq data on murine retina development.** UMAP representations of scCoGAPS patterns colored by individual pattern weights.

**Figure S3. Heatmap of FDR corrected log10 pvals for GO enrichment in scCoGAPS patterns.** All GO pathways with p-values <.01 after BH correction were included.

**Figure S4. Heatmap of FDR corrected log10 pvals for KEGG enrichment in scCoGAPS patterns.** All KEGG pathways with p-values <.01 after BH correction were included.

**Figure S5. AUCs values from projected pattern weights in scCoGAPS patterns** (A) Heatmap of AUC values for projected murine scRNAseq of P14 mouse retina from Macosko et al. (B) Heatmap of AUC values for projected human bulk RNA-Seq time course of the developing retina from Hoshino et al.

**Figure S6. ATAC-Seq from Chx10-GFP-positive murine retinal progenitor cell time-course analysis projected into scCoGAPS 10x patterns.** Graphs indicating chromatin accessibility of scCoGAPS pattern marker genes within the RPC ATAC-seq datasets.

**Figure S7. Alluvial plots of brain projections.** Alluvial plot of projected patterns link previously annotated cell types to scCoGAPS patterns for which at least 75% (A+C) or 50% (B) of cells of a given type have a significant projection (Wald test; BH-correction; q < .01) in scRNAseq from human cortical development from (A) Nowakowski, et al and (B) Zhong, et al. as well as mouse midbrain development from (C) La Manno, et al.

**Figure S8. AUC heatmaps for developing brain scRNA-Seq in scCoGAPS patterns of retina development.** Heatmaps of AUC values for scRNAseq from (A) Nowakowski, et al, (B) Zhong, et al. and (C) La Manno et al projected into 80 scCoGAPS patterns of murine retina development.

**Figure S9. Tabular Muris projection supplement.** (A) Alluvial plot of projected patterns links previously annotated cell types to scCoGAPS patterns for which at least 50% of cell of a given type have a significant projection (Wald test; BH-correction; q < .01) in scRNAseq from the Tabula Muris dataset (B) UMAP representation of scRNA-Seq data from the Tabular Muris collection colored by tissue of origin.(C) *Ins1* and (D) *Ins2* expression in a subpopulation of lung cells enriched for amacrine/horizontal cell-enriched Patterns 16 and 17. (E) Heat map showing retinal expression levels of marker genes of pattern 16 that are enriched in Ins1/2-positive lung cells.

## SUPPLEMENTAL FILES

**File S1 -** Derivation of model for CoGAPS and scCoGAPS.

## SUPPLEMENTAL TABLES

**Table S1 -** Amplitude values and pattern weights from GWCoGAPS on retina development time course of Chx:GFP positive and negative bulk RNA-Seq samples.

**Table S2 -** Amplitude values and pattern weights for scCoGAPS analysis of mouse retinal development scRNA-Seq data

**Table S3 -** PatternMarkers for scCoGAPS analysis of mouse retinal development scRNA-Seq data

**Table S4 -** Gene set enrichment of scCoGAPS amplitude values from analysis of mouse retinal development scRNA-Seq data

## REFERENCES

Aldiri, I., Xu, B., Wang, L., Chen, X., Hiler, D., Griffiths, L., Valentine, M., Shirinifard, A., Thiagarajan, S., Sablauer, A., et al. (2017). The Dynamic Epigenetic Landscape of the Retina During Development, Reprogramming, and Tumorigenesis. Neuron 94, 550–568.e10.

Bassett, E.A., and Wallace, V.A. (2012). Cell fate determination in the vertebrate retina. Trends Neurosci. 35, 565–573.

Bidaut, G., and Ochs, M.F. (2004). ClutrFree: cluster tree visualization and interpretation. Bioinformatics 20, 2869–2871.

Blackshaw, S., Fraioli, R.E., Furukawa, T., and Cepko, C.L. (2001). Comprehensive analysis of photoreceptor gene expression and the identification of candidate retinal disease genes. Cell 107, 579–589.

Blackshaw, S., Harpavat, S., Trimarchi, J., Cai, L., Huang, H., Kuo, W.P., Weber, G., Lee, K., Fraioli, R.E., Cho, S.-H., et al. (2004). Genomic analysis of mouse retinal development. PLoS Biol. 2, E247.

Brunet, J.-P., Tamayo, P., Golub, T.R., and Mesirov, J.P. (2004). Meta-genes and molecular pattern discovery using matrix factorization. Proceedings of the National Academy of Sciences 101, 4164–4169.

Buenrostro, J.D., Giresi, P.G., Zaba, L.C., Chang, H.Y., and Greenleaf, W.J. (2013). Transposition of native chromatin for fast and sensitive epigenomic profiling of open chromatin, DNA-binding proteins and nucleosome position. Nat. Methods 10, 1213–1218.

Butler, A., and Satija, R. (2017). Integrated analysis of single cell transcriptomic data across conditions, technologies, and species. bioRxiv.

Ching, T., Huang, S., and Garmire, L.X. (2014). Power analysis and sample size estimation for RNA-Seq differential expression. RNA 20, 1684–1696.

Clark, B., Stein-O’Brien, G., Shiau, F., Cannon, G., Davis, E., Sherman, T., Rajaii, F., James-Esposito, R., Gronostajski, R., Fertig, E., et al. (2018). Comprehensive analysis of retinal development at single cell resolution identifies NFI factors as essential for mitotic exit and specification of late-born cells.

Cleary, B., Cong, L., Lander, E., and Regev, A. (2017a). Composite measurements and molecular compressed sensing for highly efficient transcriptomics. bioRxiv.

Cleary, B., Cong, L., Cheung, A., Lander, E.S., and Regev, A. (2017b). Efficient Generation of Transcriptomic Profiles by Random Composite Measurements. Cell 171, 1424–1436.e18.

Curcio, C.A., and Allen, K.A. (1990). Topography of ganglion cells in human retina. J. Comp. Neurol. 300, 5–25.

DeTomaso, D., and Yosef, N. (2016). FastProject: a tool for low-dimensional analysis of single-cell RNA-Seq data. BMC Bioinformatics 17, 315.

Eraslan, G., Simon, L.M., Mircea, M., Mueller, N.S., and Theis, F.J. (2018). Single cell RNA-seq denoising using a deep count autoencoder. bioRxiv.

Fertig, E.J., Ding, J., Favorov, A.V., Parmigiani, G., and Ochs, M.F. (2010). CoGAPS: an R/C++ package to identify patterns and biological process activity in transcriptomic data. Bioinformatics 26, 2792–2793.

Fertig, E.J., Markovic, A., Danilova, L.V., Gaykalova, D.A., Cope, L., Chung, C.H., Ochs, M.F., and Califano, J.A. (2013a). Preferential Activation of the Hedgehog Pathway by Epigenetic Modulations in HPV Negative HNSCC Identified with Meta-Pathway Analysis. PLoS One 8, e78127.

Fertig, E.J., Favorov, A.V., and Ochs, M.F. (2013b). Identifying context-specific transcription factor targets from prior knowledge and gene expression data. IEEE Trans. Nanobioscience 12, 142–149.

Hendrickson, A., and Drucker, D. (1992). The development of parafoveal and mid-peripheral human retina. Behav. Brain Res. 49, 21–31.

Hendrickson, A., Possin, D., Vajzovic, L., and Toth, C.A. (2012). Histologic development of the human fovea from midgestation to maturity. Am. J. Ophthalmol. 154, 767–778.e2.

Hicks, S.C., Townes, F.W., Teng, M., and Irizarry, R.A. (2017). Missing data and technical variability in single-cell RNA-sequencing experiments. Biostatistics.

Hoshino, A., Ratnapriya, R., Brooks, M.J., Chaitankar, V., Wilken, M.S., Zhang, C., Starostik, M.R., Gieser, L., La Torre, A., Nishio, M., et al. (2017). Molecular Anatomy of the Developing Human Retina. Dev. Cell 43, 763–779.e4.

Javed, A., and Cayouette, M. (2017). Temporal Progression of Retinal Progenitor Cell Identity: Implications in Cell Replacement Therapies. Front. Neural Circuits 11, 105.

Keren-Shaul, H., Spinrad, A., Weiner, A., Matcovitch-Natan, O., Dvir-Szternfeld, R., Ulland, T.K., David, E., Baruch, K., Lara-Astaiso, D., Toth, B., et al. (2017). A Unique Microglia Type Associated with Restricting Development of Alzheimer’s Disease. Cell 169, 1276–1290.e17.

Kim, D., Langmead, B., and Salzberg, S.L. (2015). HISAT: a fast spliced aligner with low memory requirements. Nat. Methods 12, 357–360.

Kim, D., Langmead, B., and Salzberg, S.L. (2016). HISAT2 implementation.

Kiselev, V.Y., Yiu, A., and Hemberg, M. (2018). scmap: projection of single-cell RNA-seq data across data sets. Nat. Methods 15, 359–362.

Kohwi, M., and Doe, C.Q. (2013). Temporal fate specification and neural progenitor competence during development. Nat. Rev. Neurosci. 14, 823–838.

Kossenkov, A.V., Peterson, A.J., and Ochs, M.F. (2007). Determining transcription factor activity from microarray data using Bayesian Markov chain Monte Carlo sampling. Stud. Health Technol. Inform. 129, 1250–1254.

La Manno, G., Gyllborg, D., Codeluppi, S., Nishimura, K., Salto, C., Zeisel, A., Borm, L.E., Stott, S.R.W., Toledo, E.M., Villaescusa, J.C., et al. (2016). Molecular Diversity of Midbrain Development in Mouse, Human, and Stem Cells. Cell 167, 566–580.e19.

Langmead, B., and Salzberg, S.L. (2012). Fast gapped-read alignment with Bowtie 2. Nat. Methods 9, 357–359.

Lee, D.D., and Seung, H.S. (1999). Learning the parts of objects by non-negative matrix factorization. Nature.

Leek, J.T., Johnson, W.E., Parker, H.S., Jaffe, A.E., and Storey, J.D. (2012). The sva package for removing batch effects and other unwanted variation in high-throughput experiments. Bioinformatics 28, 882–883.

Li, Y., and Ngom, A. (2013). The non-negative matrix factorization toolbox for biological data mining. Source Code Biol. Med. 8, 10.

Li, H., Handsaker, B., Wysoker, A., Fennell, T., Ruan, J., Homer, N., Marth, G., Abecasis, G., Durbin, R., and 1000 Genome Project Data Processing Subgroup (2009). The Sequence Alignment/Map format and SAMtools. Bioinformatics 25, 2078–2079.

Liao, Y., Smyth, G.K., and Shi, W. (2014). featureCounts: an efficient general purpose program for assigning sequence reads to genomic features. Bioinformatics 30, 923–930.

Loos, C., Moeller, K., Fröhlich, F., Hucho, T., and Hasenauer, J. (2018). A Hierarchical, Data-Driven Approach to Modeling Single-Cell Populations Predicts Latent Causes of Cell-To-Cell Variability. Cell Syst 6, 593–603.e13.

Macosko, E.Z., Basu, A., Satija, R., Nemesh, J., Shekhar, K., Goldman, M., Tirosh, I., Bialas, A.R., Kamitaki, N., Martersteck, E.M., et al. (2015). Highly Parallel Genome-wide Expression Profiling of Individual Cells Using Nanoliter Droplets. Cell 161, 1202–1214.

Miller, F.D., and Gauthier, A.S. (2007). Timing is everything: making neurons versus glia in the developing cortex. Neuron 54, 357–369.

Moloshok, T.D., Klevecz, R.R., Grant, J.D., Manion, F.J., Speier, W.F., 4th, and Ochs, M.F. (2002). Application of Bayesian decomposition for analysing microarray data. Bioinformatics 18, 566–575.

Nowakowski, T.J., Bhaduri, A., Pollen, A.A., Alvarado, B., Mostajo-Radji, M.A., Di Lullo, E., Haeussler, M., Sandoval-Espinosa, C., Liu, S.J., Velmeshev, D., et al. (2017). Spatiotemporal gene expression trajectories reveal developmental hierarchies of the human cortex. Science 358, 1318–1323.

O’Brien, K.M.B., Schulte, D., and Hendrickson, A.E. (2003). Expression of photoreceptor-associated molecules during human fetal eye development. Mol. Vis. 9, 401–409.

Ochs, M.F., and Fertig, E.J. (2012). Matrix Factorization for Transcriptional Regulatory Network Inference. … Bioinformatics and Computational Biology … 1–10.

Ochs, M.F., Rink, L., Tarn, C., Mburu, S., Taguchi, T., Eisenberg, B., and Godwin, A.K. (2009). Detection of Treatment-Induced Changes in Signaling Pathways in Gastrointestinal Stromal Tumors Using Transcriptomic Data. Cancer Res. 69, 9125–9132.

Pan, S.J., Kwok, J.T., and Yang, Q. (2008). Transfer learning via dimensionality reduction. AAAI.

Quinlan, A.R., and Hall, I.M. (2010). BEDTools: a flexible suite of utilities for comparing genomic features. Bioinformatics 26, 841–842.

Ritchie, M.E., Phipson, B., Wu, D., Hu, Y., Law, C.W., Shi, W., and Smyth, G.K. (2015). limma powers differential expression analyses for RNA-sequencing and microarray studies. Nucleic Acids Res. 43, e47.

Rowan, S., and Cepko, C.L. (2004). Genetic analysis of the homeodomain transcription factor Chx10 in the retina using a novel multifunctional BAC transgenic mouse reporter. Dev. Biol. 271, 388–402.

Rozenblatt-Rosen, O., Stubbington, M.J.T., Regev, A., and Teichmann, S.A. (2017). The Human Cell Atlas: from vision to reality. Nature 550, 451–453.

Sibisi, S., and Skilling, J. (1996). Bayesian Density Estimation. In Maximum Entropy and Bayesian Methods, pp. 189–198.

Sing, T., Sander, O., Beerenwinkel, N., and Lengauer, T. (2005). ROCR: visualizing classifier performance in R. Bioinformatics 21, 3940–3941.

Skilling, J., and Sibisi, S. (1996). Priors on Measures. In Maximum Entropy and Bayesian Methods, pp. 261–270.

Stein-O’Brien, G.L., Arora, R., Culhane, A.C., Favorov, A., Greene, C., Goff, L.A., Li, Y., Ngom, A., Ochs, M.F., Xu, Y., et al. (2017a). Enter the matrix: Interpreting unsupervised feature learning with matrix decomposition to discover hidden knowledge in high-throughput omics data.

Stein-O’Brien, G.L., Carey, J.L., Lee, W.-S., Considine, M., Favorov, A.V., Flam, E., Guo, T., Li, S., Marchionni, L., Sherman, T., et al. (2017b). PatternMarkers & GWCoGAPS for novel data-driven biomarkers via whole transcriptome NMF. Bioinformatics.

Tasic, B., Menon, V., Nguyen, T.N., Kim, T.K., Jarsky, T., Yao, Z., Levi, B., Gray, L.T., Sorensen, S.A., Dolbeare, T., et al. (2016). Adult mouse cortical cell taxonomy revealed by single cell transcriptomics. Nat. Neurosci. 19, 335.

Torrey, L., and Shavlik, J. (2009). Transfer Learning. In Handbook of Research on Machine Learning Applications and Trends Algorithms, Methods, and Techniques, E.S. Olivas, ed. pp. 242–264.

Trapnell, C., Hendrickson, D.G., Sauvageau, M., Goff, L., Rinn, J.L., and Pachter, L. (2012). Differential analysis of gene regulation at transcript resolution with RNA-seq. Nat. Biotechnol. 31, 46–53.

Tung, P.-Y., Blischak, J.D., Hsiao, C.J., Knowles, D.A., Burnett, J.E., Pritchard, J.K., and Gilad, Y. (2017). Batch effects and the effective design of single-cell gene expression studies. Sci. Rep. 7, 39921.

Wagner, A., Regev, A., and Yosef, N. (2016). Revealing the vectors of cellular identity with single-cell genomics. Nat. Biotechnol.

Wang, W., Arora, R., Livescu, K., and Bilmes, J.A. (2015). Unsupervised learning of acoustic features via deep canonical correlation analysis. In 2015 IEEE International Conference on Acoustics, Speech and Signal Processing (ICASSP), pp. 4590–4594.

Wu, A.R., Wang, J., Streets, A.M., and Huang, Y. (2017). Single-Cell Transcriptional Analysis. Annu. Rev. Anal. Chem. 10, 439–462.

Wysoker, A., Tibbetts, K., and Fennell, T. (2013). Picard tools version 1.90.

Wyss-Coray, T., Darmanis, S., and Muris Consortium, T. (2018). Single-cell transcriptomic characterization of 20 organs and tissues from individual mice creates a Tabula Muris. bioRxiv.

Young, R.W. (1984). Cell death during differentiation of the retina in the mouse. J. Comp. Neurol. 229, 362–373.

Zhong, S., Zhang, S., Fan, X., Wu, Q., Yan, L., Dong, J., Zhang, H., Li, L., Sun, L., Pan, N., et al. (2018). A single-cell RNA-seq survey of the developmental landscape of the human prefrontal cortex. Nature.

Zhu, X., Ching, T., Pan, X., Weissman, S.M., and Garmire, L. (2017). Detecting heterogeneity in single-cell RNA-Seq data by nonnegative matrix factorization. PeerJ 5, e2888.

Zibetti C., Liu S., Wan J. Qian J., and Blackshaw S. (2017) Lhx2 regulates temporal changes in chromatin accessibility and transcription factor binding in retinal progenitor cells. BioRxiv. https://doi.org/10.1101/238279

